# Trivalent NDV-HXP-S vaccine protects against phylogenetically distant SARS-CoV-2 variants of concern in mice

**DOI:** 10.1101/2022.03.21.485247

**Authors:** Irene González-Domínguez, Jose Luis Martínez, Stefan Slamanig, Nicholas Lemus, Yonghong Liu, Tsoi Ying Lai, Juan Manuel Carreño, Gagandeep Singh, Gagandeep Singh, Michael Schotsaert, Ignacio Mena, Stephen McCroskery, Lynda Coughlan, Florian Krammer, Adolfo García-Sastre, Peter Palese, Weina Sun

## Abstract

Equitable access to vaccines is necessary to limit the global impact of the coronavirus disease 2019 (COVID-19) pandemic and the emergence of new severe acute respiratory syndrome coronavirus 2 (SARS-CoV-2) variants. In previous studies, we described the development of a low-cost vaccine based on a Newcastle Disease virus (NDV) expressing the prefusion stabilized spike protein from SARS-CoV-2, named NDV-HXP-S. Here, we present the development of next-generation NDV-HXP-S variant vaccines, which express the stabilized spike protein of the Beta, Gamma and Delta variants of concerns (VOC). Combinations of variant vaccines in bivalent, trivalent and tetravalent formulations were tested for immunogenicity and protection in mice. We show that the trivalent preparation, composed of the ancestral Wuhan, Beta and Delta vaccines, substantially increases the levels of protection and of cross-neutralizing antibodies against mismatched, phylogenetically distant variants, including the currently circulating Omicron variant.

## Introduction

Severe acute respiratory syndrome coronavirus 2 (SARS-CoV-2) is the causative agent of the current coronavirus disease 2019 (COVID-19). Since the beginning of the pandemic, the emergence of new variants of concern (VOC) has threatened the protection conferred by vaccines based on the original strain (Figure 1) [1]. In December 2020, the Alpha variant (B.1.1.7) and Beta variant (B.1.351) were declared VOC and spread over the world, followed by the Gamma strain (P.1) that was declared a VOC in January 2021. Both Beta and Gamma variants exhibited notable resistance to neutralizing antibodies raised against the original strain in humans [1, 2]. In May 2021, a strong pandemic wave in India gave rise to a new VOC: the Delta variant (B.1.617.2). This new VOC harbored different mutations in the spike that significantly reduced its sensitivity to neutralizing antibodies, and increased transmissibility quickly replacing the previous variants worldwide (Figure 1B) [1, 3]. In November 2021, a new VOC named Omicron appeared in South Africa. Since then, Omicron has taken over worldwide, replacing the Delta variant [4]. Compared to the previous VOC, Omicron presents the highest number of mutations in the spike protein and has shown the highest drop-in neutralization activity [5, 6]. Currently, the Omicron sub-lineage BA.2, seems to show even more immune evasion and transmissibility [7–9].

**Figure 1:**
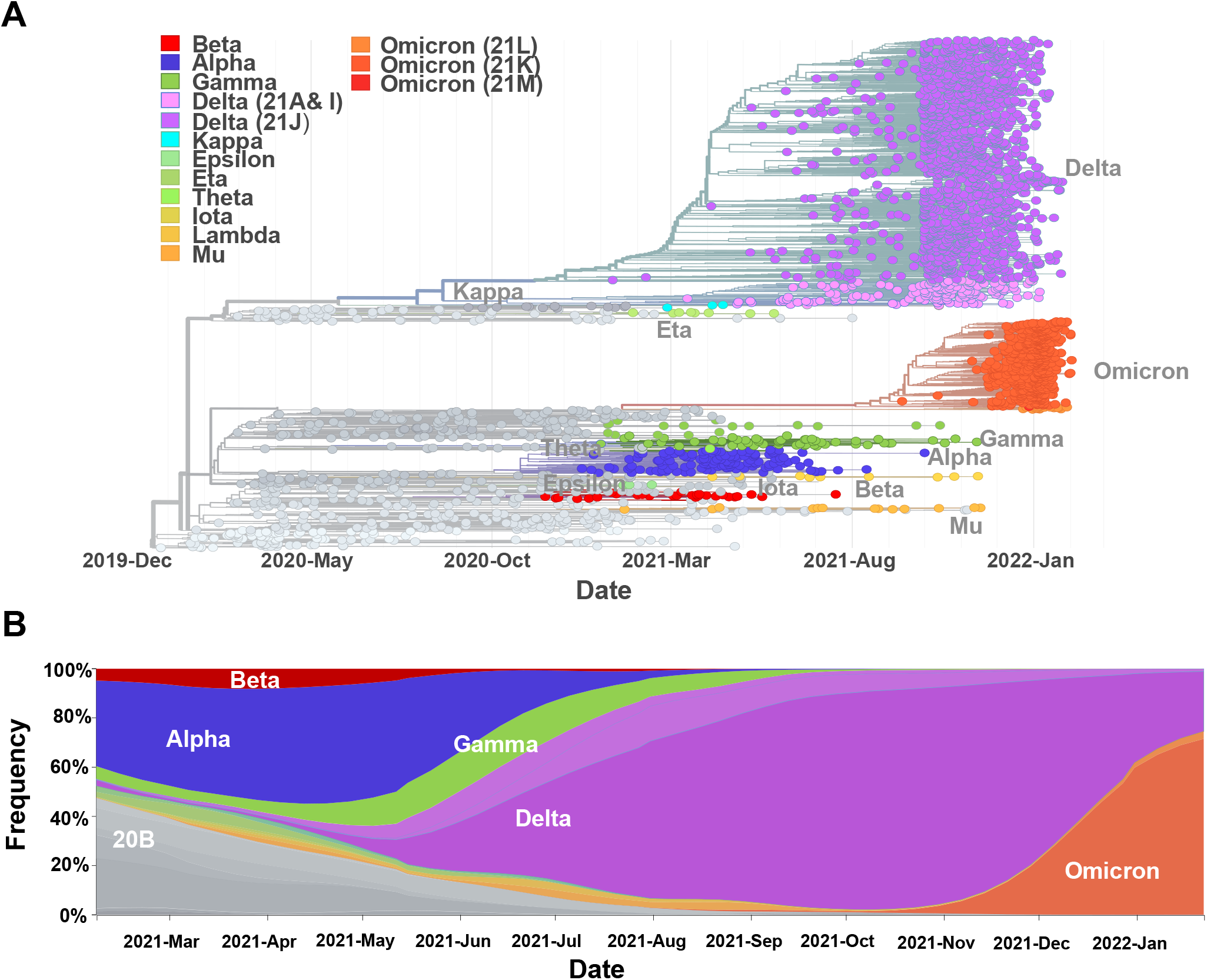
Evolution of SARS-CoV-2 and appearance of variants of concern (VOC) **(A)** Phylogenetic tree (15-Dec-2019 to 06-Feb-2022) with 3057 genomes showing the global evolutionary relationships of SARS-CoV-2 viruses from the ongoing COVID-19 pandemic **(B)** Timeline (15-Dec-2019 to 06-Feb-2022) graph showing the global frequencies by clade of the different SARS-CoV-2 viruses. Graphics were adapted from the website /nextstrain.org/ncov/gisaid/global (accessed 05-Feb-2022 CC-BY [29, 30])

Despite the unprecedented, rapid development of COVID-19 vaccines, only ~63% of the global population are fully vaccinated (as of 14^th^ March 2022) [6]. Hence, there is still a need for COVID-19 vaccines that can be produced locally in low- and middle-income countries (LMICs), where the vaccination rates are the lowest worldwide [6]. In previous work, we developed a vaccine candidate named NDV-HXP-S [10] that can be manufactured like influenza virus vaccines at low cost in embryonated chicken eggs in facilities located globally [11]. This vaccine is based on an avirulent Newcastle disease virus (NDV) strain which presents a SARS-CoV-2 spike protein stabilized in its prefusion-conformation by the introduction of six proline mutations (HexaPro, HXP-S) [4, 12] and elimination of the furin cleavage site. NDV-HXP-S can be used as live vaccine [11, 13, 14] or as an inactivated vaccine [11, 15]. Clinical trials with a live version are ongoing in Mexico (NCT04871737) and the US (NCT05181709), while the inactivated vaccine is being tested in Vietnam (NCT04830800), Thailand (NCT04764422) and Brazil (NCT04993209). Interim results from the initial Phase I/II trials have demonstrated that the vaccine was safe and immunogenic [15–17].

Here, we present the development of NDV-HXP-S variant vaccines displaying the VOC spike proteins of Beta, Gamma and Delta. We tested the immunogenicity and protection induced by vaccination with inactivated NDV-HXP-S variants in mice. We observed that variant-specific vaccines induced the strongest antibody responses towards the homologous SARS-CoV-2 VOC. Furthermore, we found that a combination of multivalent NDV-HXP-S with the Wuhan, Beta and Delta provided a broader protection against a panel of VOC than the monovalent formulations.

## Results

### Design and production of NDV-HXP-S variant vaccines

NDV-HXP-S variant vaccines based on three VOC, Beta (B.1.351), Gamma (P.1) and Delta (B.1.617.2), were rescued using reverse genetics in mammalian cell cultures and further amplified in specific-pathogen free (SPF) embryonated chicken eggs as previously described (Figure 2) [11]. The nucleotide sequence of the constructs was codon-optimized for mammalian host expression. The HXP-S sequence was inserted between the P and M genes of the NDV genome. We removed the polybasic cleavage site (^682^RRAR^685^) and replaced the transmembrane domain (TM) and cytoplasmic tail (CT) of the spike with those from the fusion (F) protein of La Sota NDV. The HXP mutations were introduced into the spike S2 region of all three variant S proteins to improve their stability as reported previously (Figure 2A & 2B) [12]. In addition, we found that a completely cleaved Delta spike was observed when the Delta specific S-P681R mutation was included. Therefore, we kept the 681 as proline to ensure the homogeneity of prefusion conformation (S0) (Figure 2C).

**Figure 2:**
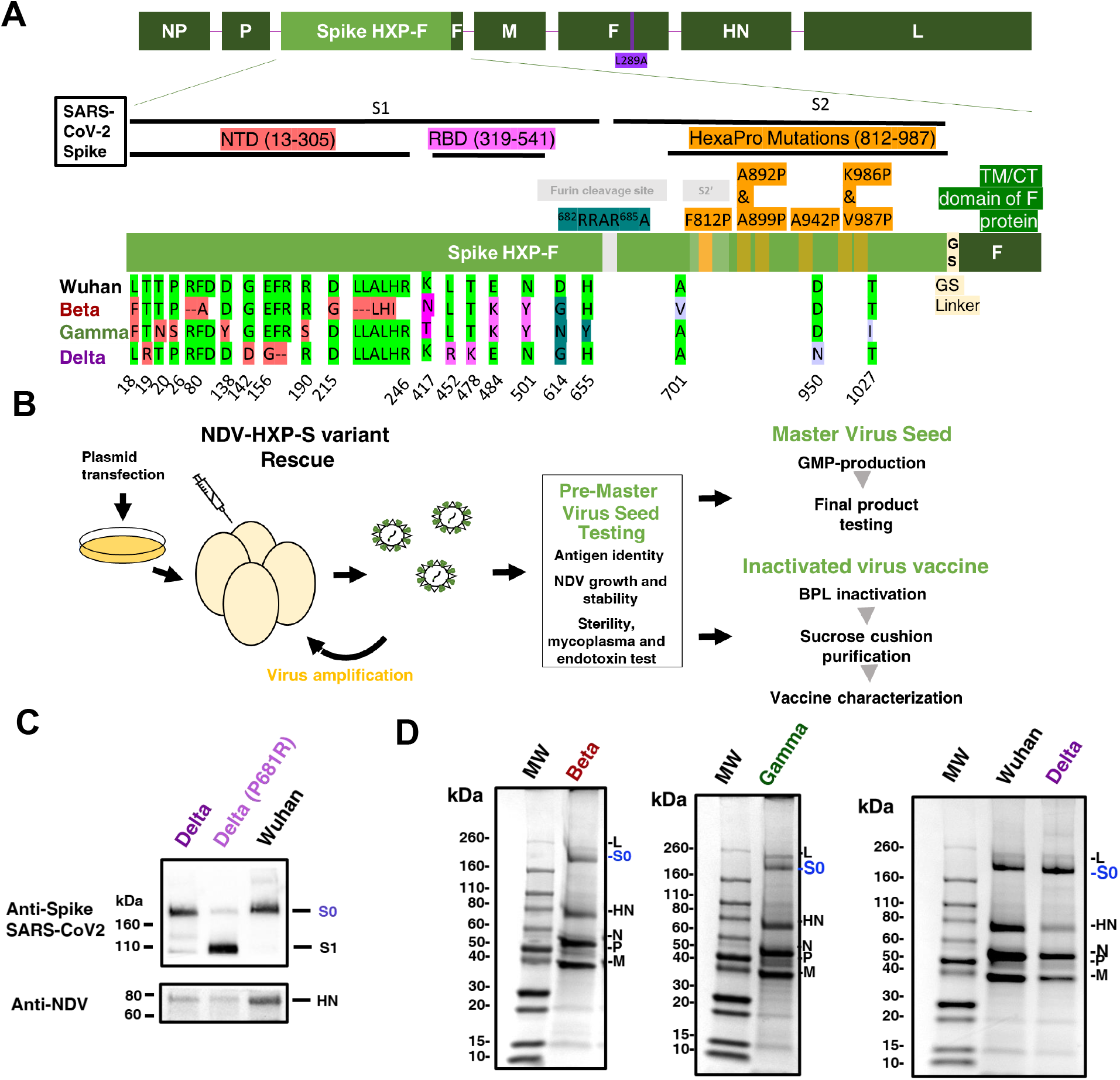
Design, production, and characterization of NDV-HXP-S variant vaccines. **(A)** Structure and design of the NDV-HXP-S construct. The different SARS-CoV-2 spike sequences were introduced between the P and M genes of LaSota L289A NDV strain. The ectodomain of the spike was connected to the transmembrane domain and cytoplasmic tail (TM/CT) of the F protein of the NDV. The original polybasic cleavage site was removed by mutating RRAR to A. The HexaPro (F817P, A892P, A899P, A942P, K986P and V987P) stabilizing mutations were introduced. The sequence was codon-optimized for mammalian host expression. Original WuhanHXP-S sequence is aligned with the new variant constructs: Beta, Gamma and Delta. Changes in the N-terminal domain (NTD), the Receptor Binding Domain (RBD) and in the Spike 2 (S2) with respect to the original WuhanHXP-S sequence are highlighted in red, pink and light purple, respectively. **(B)** NDV-HXP-S variants were rescued by reverse genetics as previously described [24]. Cells were co-transfected with the expression plasmid required for replication and transcription of the NDV viral genome (NP, P, and L), together with the full length NDV cDNA. After 2 or 3 days, the tissue culture supernatants were inoculated into eight- or nine-day-old specific pathogen free (SPF) embryonated chicken eggs. Antigen identity was confirmed by biochemical methods and sequencing. The genetic stability of the recombinant viruses was evaluated across multiples passages on ten days old-SPF embryonated chicken eggs. NDV-HXP-S vaccine was inactivated with BPL and purified by sucrose cushion ultracentrifugation **(C)** Comparison of NDV-HXP-S Delta virus versus NDV-HXP-S Delta with P681R mutation. The spike protein and NDV HN proteins were detected by western blot using an anti-spike 2B3E5 mouse monoclonal antibody and an anti-HN 8H2 mouse monoclonal antibody, respectively. **(D)** Protein staining of NDV-HXP-S variant vaccines resolved on 4-20% SDS-PAGE. The viral proteins were visualized by Coomassie Blue staining (L, S0, HN, N, P and M). The uncleaved S0 spike protein is highlighted in blue with an approximate size of 200 kDa.

Research-grade beta-propiolactone (BPL) inactivated NDV-HXP-S variant vaccine preparations were produced. Virus was concentrated from the harvested allantoic fluid through a sucrose cushion and resolved on a sodium dodecyl–sulfate polyacrylamide gel (SDS–PAGE) with Coomassie Blue staining to evaluate the presence of all NDV-HXP-S proteins. NDV-HXP-S variants showed an extra band between 160 kDa and 260 kDa below the L protein of the NDV that corresponds to the size of the uncleaved spike (S0, Figure 2D) [11].

### NDV-HXP-S variant vaccines protect against homologous challenge in mouse models

The immunogenicity and *in vivo* protection of the new NDV-HXP-S Beta and Gamma vaccine candidates was first evaluated in BALB/c mice transduced with a non-replicating human adenovirus 5 expressing human angiotensin-converting enzyme 2 (Ad5-hACE2) (Figure 3A). A two-dose intramuscular (IM) vaccination regimen with a total protein content of 1 μg per mouse of purified NDV-HXP-S vaccine candidate with a 3-week period between doses was followed. Three weeks after the boost, BALB/c mice were transduced by intranasal (IN) administration of the Ad5-hACE2 and five days later mice were challenged with SARS-CoV-2 viruses via the IN route, as previously described [18]. Two days after challenge, viral titers in the lung homogenates were quantified by plaque assays. Vaccination with wild type NDV was used as negative control (Figure 3B).

**Figure 3:**
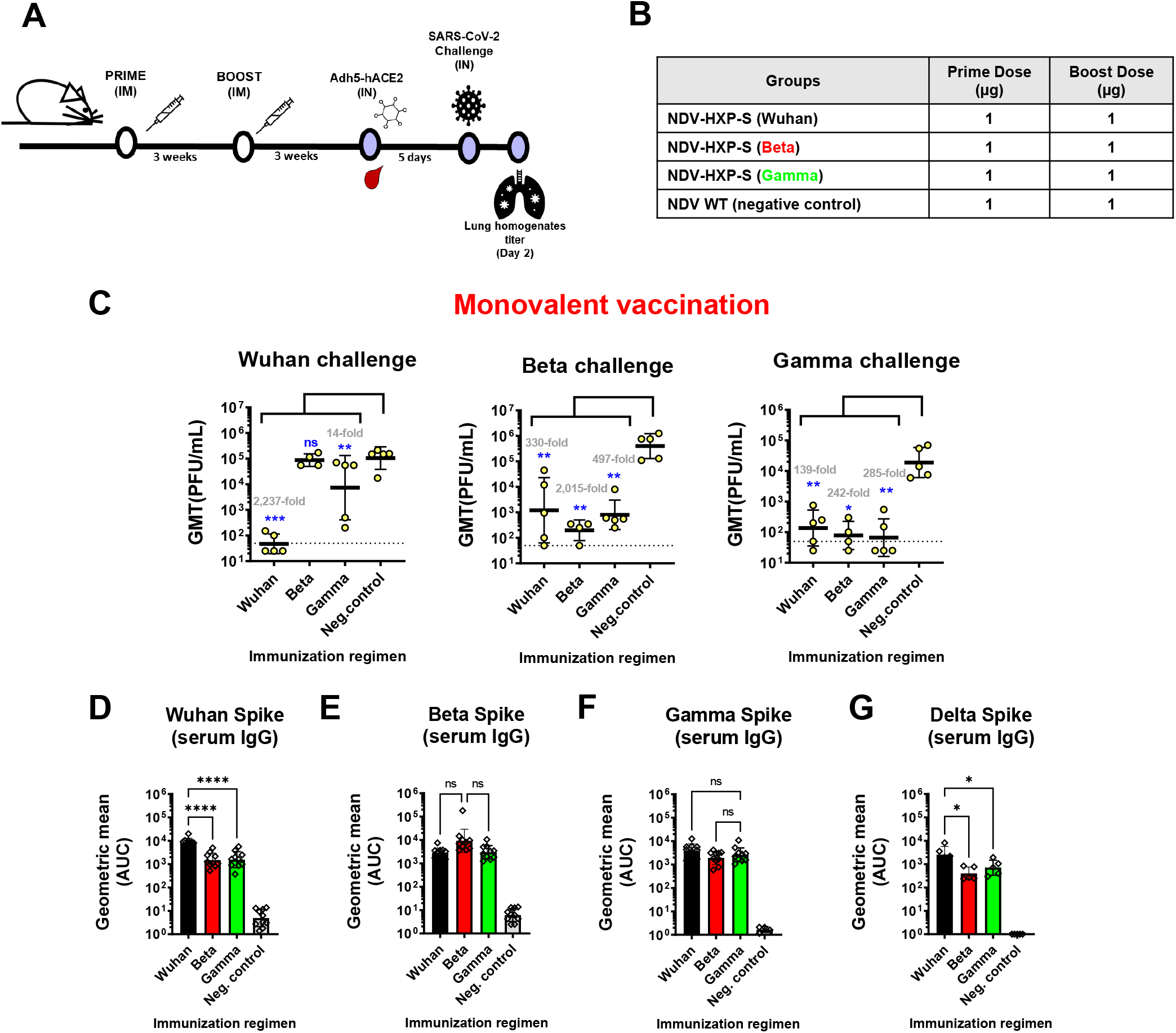
NDV-HXP-S Beta and Gamma induce protective antibodies against homologous infection. **(A&B)** Design of the study and groups. Eight-to ten-week-old female BALB/c mice were used either vaccinated with 1 μg of total dose of inactivated NDV-HXP-S variant vaccines or WT NDV (negative control). Two immunizations were performed via the intramuscular route (IM) at D0 and D21. At D44, mice were treated with Ad5-hACE2. At D49, mice were challenged with a Wuhan-like (USA-WA1/2020), Beta (B.1.351) or Gamma (P.1) strains and at day two after challenge, lungs were harvested and homogenized in 1 mL PBS and titers were measured by plaque assay on Vero E6 cells. Viral titers in lung homogenates after Wuhan, Beta or Gamma challenge (n=5) **(C).** Viral titers were measured by plaque assay on Vero E6 cells and plotted as Geometric mean titer (GMT) of PFU/mL (limit of detection equals to 50 PFU/mL; a titer of 25 PFU/mL was assigned to negative samples). The error bars represent geometric standard deviation. Geometric mean fold titers over the control are indicated in gray. Spike-specific serum IgG levels against Wuhan spike (n=10) **(D)**, Beta spike (n=10) **(E)**, Gamma spike (n=10) **(F)** and Delta spike proteins (n=5) **(G)**. Antibodies in post-boost (D43) sera samples from the different immunization regimens were measured by ELISA. The GMT of the area under the curve (AUC) were graphed. The error bars represent geometric standard deviation. Statistical difference was analyzed by ordinary one-way ANOVA corrected for Dunnett’s multiple comparisons test in all figures (*p < 0.05; **p < 0.01; ***p < 0.001; ****p < 0.0001).

A viral titer reduction of 2,237-fold, 2,015-fold and 285-fold was obtained in the homologous challenge using monovalent vaccination regimens with Wuhan, Beta and Gamma NDV-HXP-S vaccines compared to the negative control, respectively (*p.value <0.05 versus the neg.control*, Figure 3C). Heterologous protection with Wuhan NDV-HXP-S against Beta and Gamma challenge was observed with a viral titer reduction of 51-fold and 120-fold compared to the negative control, respectively, in agreement with previous studies [11]. Beta NDV-HXP-S vaccine showed cross-protection against the Gamma variant challenge, and *vice versa*. However, an asymmetric weaker protection of Beta and Gamma vaccines was observed against the Wuhan challenge.

Together with the *in vivo* challenge study, mice were bled 3-weeks after boost to measure the spike-specific IgG levels. Antibody titers after vaccination against Wuhan, Beta and Gamma spikes were compared in Figures 3D-F, respectively. Wuhan spike-specific IgG serum antibody titers were higher after the vaccination with the original construct compared to Beta and Gamma NDV-HXP-S vaccinations (Figure 3D). Similar Beta and Gamma spike-specific antibody titers were obtained with all three vaccination regimens (Figure 3E-F). These results correlate with the challenge against the three viruses, where a similar cross-protection was obtained against Beta and Gamma challenges with all three vaccination regimens. As the Delta variant emerged, we also measured post-boost serum antibodies against the Delta spike (Figure 3G). Compared to the other spike-specific titers, the levels of cross-reactive antibodies against the Delta spike were reduced in all three vaccination groups, suggesting a higher drop in protection against this variant with the current constructs tested.

### Trivalent and tetravalent NDV-HXP-S variant vaccinations increase protection *in vivo* against phylogenetically distant SARS-CoV-2 variants

Following the characterizations of Beta and Gamma NDV-HXP-S vaccines, the Delta VOC appeared in India and rapidly replaced the other variants (Figure 1B). When the Delta vaccine construct was generated, a second mouse immunization and challenge study was performed testing vaccine formulations containing the Delta component (Figure 4A). Based on the previous results, the following groups were evaluated: monovalent Wuhan, monovalent Delta, bivalent Wuhan and Delta, sequential vaccination with a first dose of Wuhan followed by Delta, trivalent (Wuhan, Delta and Beta) and tetravalent (Wuhan, Delta, Beta and Gamma). In all cases, the vaccination dose was 1 μg of total protein (Figure 4B).

**Figure 4:**
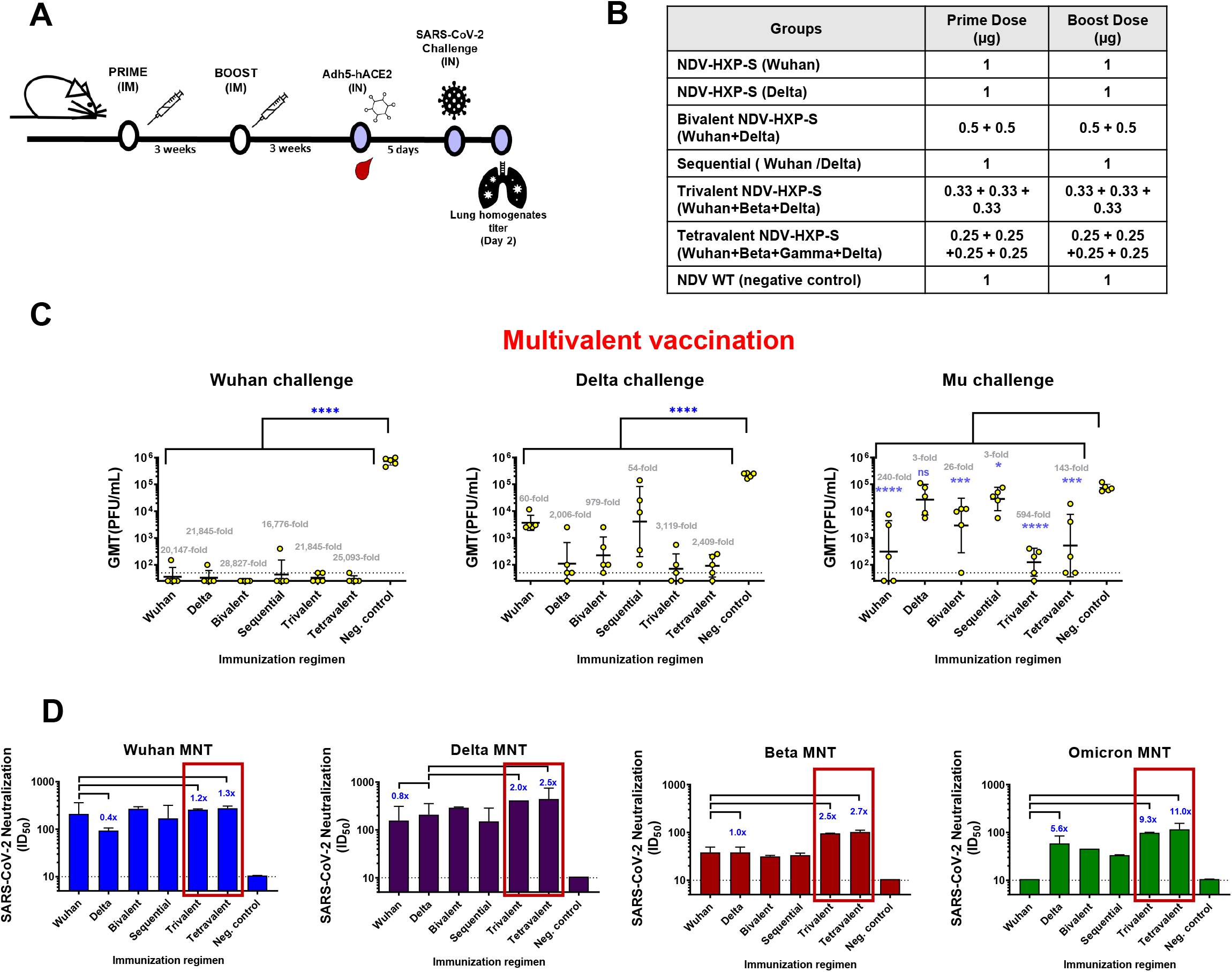
Trivalent and tetravalent NDV-HXP-S vaccination regimens induce protection against phylogenetically distant SARS-CoV-2 variants. **(A&B)** Design of the study and groups. Eight-to ten-week-old female BALB/c mice were used either vaccinated with 1 μg of total dose of inactivated NDV-HXP-S variant vaccines or WT NDV (negative control). Two immunizations were performed via the intramuscular route (IM) at D0 and D21. At D44, mice were treated with Ad5-hACE2. At D49, mice were challenged with Wuhan-like (USA-WA1/2020), Delta (B.1.617.2) or Mu (B.1.621) strains and at day two after challenge, lungs were harvested and homogenized in 1 mL PBS and titers were measured by plaque assay. **(C)** Viral titers after challenge (n=5). Viral titers were measured by plaque assay on Vero E6 cells for Wuhan-like USA-WA1/2020 challenge and Vero-TMPRSS2 cells for Delta and Mu challenges and plotted as GMT of PFU/mL (limit of detection equals to 50 PFU/mL; a titer of 25 PFU/mL was assigned to negative samples). The error bars represent geometric standard deviation. Statistical differences were analyzed by ordinary one-way ANOVA corrected for Dunnett’s multiple comparisons test. The *p values* and geometric mean fold titers over the control are indicated in blue and gray, respectively. (*p < 0.05; **p < 0.01; ***p < 0.001; ****p < 0.0001) **(D)** Panel of neutralizing activity. Post-boost pooled sera were tested in microneutralization (MNT) assays against Wuhan-like (USA-WA/2020) strain, Delta (B.1.617.2) variant, Beta (B.1.351) variant and Omicron (B.1.1.529) variant in technical duplicates. GMT serum dilutions inhibiting 50% of the infection (ID50) were plotted (limit of detection equals to 10 and was assigned to negative samples). Geometric mean fold change is added in blue.

In this study, we wanted to test not only the protection against homologous challenge, but also against a phylogenetically distant SARS-CoV-2 variant, which is unmatched to any vaccine component. To do so, animals in each group were divided into 3 subgroups and challenged with Wuhan, Delta and Mu variants (Figure 4C). As expected, Wuhan and Delta showed protection against its homologous virus challenge. Of note, bivalent, trivalent and tetravalent vaccines showed comparable levels of protection as the homologous monovalent vaccines against Washington 1 (Wuhan-like) and Delta challenges. Viral titers were reduced in trivalent condition 21,845-fold and 3,119-fold compared to the negative control in Wuhan and Delta challenges, respectively. However, sequential vaccination regimen was shown to be less protective, with only a 54-fold mean titer reduction in Delta challenge. In the case of the Mu challenge, trivalent formulation was proven to be the best with a titer reduction of 594-fold (Figure 4C, *p.value <0.0001 compared to the neg. control*). This reduction was 2.4 times higher than any of the other vaccination strategies tested including that of the tetravalent preparation (for discussion see below).

Considering the new emergence of the Omicron variant, we decided to compare the neutralizing activity of post-boost serum antibodies against the Wuhan, Delta, Beta and Omicron variants in authentic virus neutralization assays (Figure 4D). A similar 50% inhibitory dilution (ID50) was obtained against Wuhan in all vaccinated groups except for Delta (0.4-fold compared to Wuhan). In the case of Delta microneutralization, the neutralizing activity was increased in the trivalent and tetravalent (2.0-fold and 2.5-fold compared to the ancestral strain, respectively), whereas Wuhan alone and sequential showed a smaller neutralizing activity. A similar ID50 was obtained against Beta with monovalent, bivalent, and sequential strategies, whereas it was increased 2.5-fold and 2.7-fold in the trivalent and tetravalent groups, respectively. Little neutralizing activity was found against Omicron in the Wuhan vaccination, whereas the Delta vaccine showed a 5.6-fold increase compared to the vaccination with the monovalent Wuhan NDV-HXP-S. The neutralization titer against the Omicron was further increased by the trivalent and tetravalent vaccines (9.3-fold and 11.0-fold compared to the monovalent ancestral NDV-HXP-S vaccine group, respectively).

### Trivalent and tetravalent NDV-HXP-S vaccinations increase antibody binding to the spike of SARS-CoV-2 variants

In the attempt to elucidate the mechanism of cross-protection induced by the multivalent vaccine formulations, antibody-binding profiles were measured against a panel of SARS-CoV-2 spike proteins, including the Wuhan, Delta, Alpha, Beta, Gamma and Omicron (Figure 5A-B). Wuhan and Delta spike-specific IgG levels were comparable among all vaccination groups. In the case of Beta, an increase of 2.9-fold and 1.9-fold were observed in the trivalent and tetravalent vaccination compared to monovalent Wuhan group, respectively. An increase was also observed against Alpha (1.7-fold) and Gamma spikes (1.5-fold) in the trivalent condition (Figure 5A). Finally, vaccinated mice sera were also tested against Omicron spike. Omicron presents almost 40 different amino acid changes and deletions. A general reduction in antibody titers against Omicron spike was found in all groups compared to the other spikes tested (Figure 5A, see Y axes). Despite this drop, the addition of Delta and Beta increased spike-specific IgG titers against Omicron: 2.4-fold in the Delta vaccinated group, 5.7-fold with trivalent and 3.0 times with the tetravalent vaccinated group compared to the Wuhan vaccinated group, which correlates with the neutralization titers against Omicron (Figure 4D). Overall, the trivalent immunization regimen induced the highest titers of spike-specific antibody levels across all the variants tested (Figure 5B).

**Figure 5:**
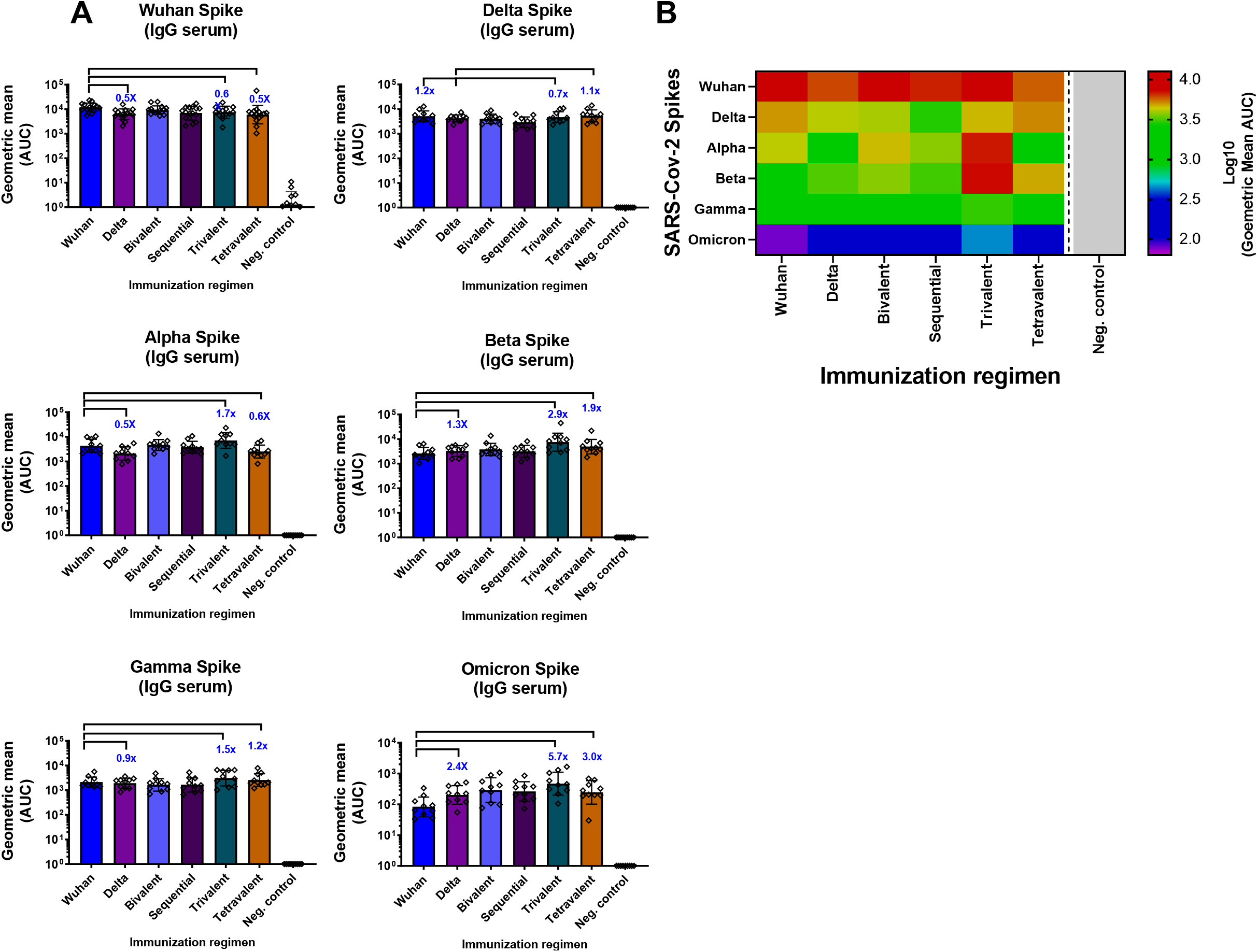
Trivalent and tetravalent NDV-HXP-S vaccination regimens induce broad serum antibody titers against phylogenetically distant SARS-CoV-2 variants. Panel **(A)** and heatmap **(B)** of spike-specific serum IgG against spike variants (n=10). Wuhan, Delta, Alpha, Beta, Gamma and Omicron spikes were used to measure the antibody binding of the different immunization groups. Antibodies in post-boost (D43) sera were measured by ELISAs. GMT AUC is depicted. The error bars represent geometric SD. Geometric mean fold change is added in blue.

## Discussion

The COVID-19 pandemic has seen an unprecedented race in the development of next generation vaccines. Compared to the traditionally slow vaccine development, novel vaccine platforms such as Moderna and Pfizer/BioNTech’s mRNA vaccines or Janssen and AstraZeneca’s adenovirus-based vaccines have been authorized for emergency use in one year [6, 19]. Despite this tremendous effort, only ~63% of the global population has been fully vaccinated, largely due to unequal vaccine distribution (as of 14^th^ of March 2022) [6]. This fact, together with the continuing emergence of VOC, which present a higher transmissibility and resistance to vaccine-mediated protection, has caused thousands of deaths per day since the beginning of the pandemic. In the last few months, the situation has worsened with the arrival of Omicron, where the world has experienced the highest number of daily cases to date [6].

With the NDV-based vaccine platform, we successfully constructed variant-specific vaccines for Beta, Gamma, and Delta variants. In mice, variant-specific vaccines effectively conferred protection against homologous viruses. To test if the co-administration of prior variants could potentially induce cross-protective immune responses against unmatched and antigenically distinct lineages, we examined bivalent, sequential, trivalent, and tetravalent formulations of inactivated NDV-HXP-S variant vaccines. Bivalent vaccination with Wuhan and Delta showed an improved immunogenicity compared to the monovalent vaccinations, whereas the sequential strategy, where the same Wuhan and Delta candidates were given in consecutive doses, showed no improvement compared to the monovalent Wuhan NDV-HXP-S group. Comparable levels of protection by a sequential administration of variant-like vaccines versus a booster dose of the original vaccine candidate have been observed in other preclinical studies [20, 21], however the long-term benefits of these heterologous boosters are still to be investigated.

Trivalent and tetravalent formulations presented synergistic responses in terms of the protection to the unmatched Mu VOC in challenge studies in mice (Figure 4C). Similarly, *in vitro* microneutralization assays have always demonstrated an increase in cross-neutralizing antibodies against all viruses tested, including against the Omicron variant (Figure 4D). Phylogenetically, Wuhan, Beta, Gamma and Delta are distinct from each other. A multivalent vaccination may induce several repertoires of cross-neutralizing antibodies that might target some common epitopes with other unmatched variants. There are several explanations to this phenomenon. One possibility is that each variant contributes with unique neutralizing epitope(s) that expand the repertoire of neutralization responses. It is also possible that conserved neutralizing epitopes shared by all the variants are more immunodominant in one variant than the other due to conformational halosteric changes resulting from spike mutations. Therefore, one variant may have induced more cross-neutralizing antibodies than the other. When trivalent and tetravalent vaccination regimens are compared between with each other, no advantage of the tetravalent regimen compared to trivalent was observed in the *in vivo* challenge experiment in mice. The dose reduction from 0.33 μg to 0.25 μg per variant or the lower immunogenicity of the Gamma vaccine might explain these results.

The NDV-HXP-S vaccine has proven its safety and immunogenicity in several preclinical and clinical studies. As an egg-based vaccine, NDV-HXP-S could be produced using the egg-based influenza virus vaccine manufacturing capacities located worldwide. Sparrow and colleagues estimated that influenza virus vaccine manufacturing capacities on its own can produce billions of doses annually [10]. Here, vaccination with a trivalent formulation in mice was performed by maintaining the total dose as that of the monovalent vaccine (1μg of total protein). Hence, the manufacturing capabilities required to produce this new formulation will remain the same.

In summary, we present the development of novel NDV-HXP-S variant vaccines and test their combination in a trivalent formulation to extend the generation of neutralizing antibodies against unmatched VOCs. This formulation exceeds the breadth of protection of the other formulations we have tested so far. This is of special interest because Omicron (B.1.1.529 or BA.1) has quickly diverged into different sub-lineages (BA.1.1, BA.2 and BA.3) and has gained prevalence globally (www.nextstrain.org/ncov). A variant-specific vaccine of any platform comes with an intrinsic disadvantage. It can only be used after the new vaccine candidate is produced and authorized by the health authorities. We believe that a more thorough selection and testing of several variant vaccines in multivalent formulations might be an effective strategy for vaccination against future SARS-CoV-2 variants.

## Data availability

Data is available upon request.

## Ethics Statement

All animal procedures in this study were performed in accordance with the animal protocol that was reviewed and approved by the Icahn School of Medicine at Mount Sinai Institutional Animal Care and Use Committee (IACUC).

## Authors Contribution

Conceptualization and design: IGD, WS, PP; NDV rescue and characterization: JLMG, SS, NL; vaccine preparation and animal experiments; IGD, YL, WS; serology: IGD, TYL, SM; protein and virus reagents; GS(a), GS(b), MS, IM, LC, AG-S; microneutralization assay: JMC, FK; data analysis: IGD, WS, PP; first draft: IGD, WS, PP; manuscript was reviewed and approved by all authors.

## Acknowledgments

We thank Dr. Benhur Lee to kindly share the BSRT7 cells, Dr. Thomas Moran for the 1C7C7 antibody and Prajakta Warang for the VERO-TMPRSS2 cell culture. We thank Dr. Randy Albrecht for support with the BSL-3 facility, procedures, and management of import/export at the Icahn School of Medicine at Mount Sinai, New York. This work was partially supported by an NIAID-funded Center of Excellence for Influenza Research and Surveillance (CEIRS, HHSN272201400008C, P.P.) and Center of Excellence for Influenza Research and Response (CEIRR, 75N93021C00014, A.G.-S., NIH grant R01 DK130425/DK/NIDDK (M.S.), NIAID R21AI157606 (L.C) and a grant from an anonymous philanthropist to Mount Sinai (P.P., F.K., A.G.-S.).

## Competing interests

The Icahn School of Medicine at Mount Sinai has filed patent applications entitled “RECOMBINANT NEWCASTLE DISEASE VIRUS EXPRESSING SARS-COV-2 SPIKE PROTEIN AND USES THEREOF” which names P.P., F.K., AG-S., and W.S. as inventors. The AG-S laboratory has received research support from Pfizer, Senhwa Biosciences, Kenall Manufacturing, Avimex, Johnson & Johnson, Dynavax, 7Hills Pharma, Pharmamar, ImmunityBio, Accurius, Hexamer, N-fold LLC, Model Medicines, Atea Pharma, Merck and Nanocomposix, and AG-S has consulting agreements for the following companies involving cash and/or stock: Vivaldi Biosciences, Contrafect, 7Hills Pharma, Avimex, Vaxalto, Pagoda, Accurius, Esperovax, Farmak, Applied Biological Laboratories, Pharmamar, Paratus, CureLab Oncology, CureLab Veterinary, Synairgen and Pfizer. The Icahn School of Medicine at Mount Sinai has filed patent applications relating to SARS-CoV-2 serological assays (U.S. Provisional Application Numbers: 62/994,252, 63/018,457, 63/020,503 and 63/024,436) which list Florian Krammer as co-inventor. Mount Sinai has spun out a company, Kantaro, to market serological tests for SARS-CoV-2. Florian Krammer has consulted for Merck and Pfizer (before 2020), and is currently consulting for Pfizer, Third Rock Ventures, Seqirus and Avimex. The Krammer laboratory is also collaborating with Pfizer on animal models of SARS-CoV-2. All other authors declared no competing interests.

## Materials and methods

### Cells

DF-1, (ATCC® CRL-12203), BSRT7 [22], and VERO-E6 (ATCC, CRL-1586) were maintained in Dulbecco’s Modified Eagle’s Medium (DMEM; Gibco, MA, USA) containing 10% (vol/vol) fetal bovine serum (FBS), 100 unit/mL of penicillin, 100 μg/mL of streptomycin (P/S; Gibco) and 10 mM 4-(2-hydroxyethyl)-1-piperazineethanesulfonic acid (HEPES) at 37°C with 5% CO_2_. VERO-TMPRSS2 cells (BPS Biosciences, #78081) were maintained in DMEM (Gibco) containing 10% (vol/vol) FBS, 100 unit/mL of penicillin, 100 μg/mL of streptomycin (P/S; Gibco), 5 mL of Nonessential Amino Acid Solution (NEAA, Corning™ MEM, NY, USA), 3 μg/mL puromycin (Invivogen, CA, USA) and 0.1 mg/mL Normocin (Invivogen) at 37°C with 5% CO_2_.

### Plasmids

Spike variant mutations were introduced into the HXP-S sequence in silico and the new constructs were obtained as synthetic double-stranded DNA fragments from Integrated DNA Technologies, using the gBlocks® Gene Fragments service and PCR [11]. Briefly, the variant HXP-S were inserted into the pNDV_LS/L289A rescue plasmid (between P and M genes) by in-Fusion cloning (Clontech, CA, USA). The recombinant product was transformed into MAX Efficiency™ Stbl2™ Competent Cells (Thermo Fisher Scientific, MA, USA) to generate the pNDV-HXP-S rescue plasmid. The plasmid was purified using PureLink™ HiPure Plasmid Maxiprep Kit (Thermo Fisher Scientific).

### Rescue of the NDV-HXP-S

As described in our previous studies [23], BSRT7 cells stably expressing the T7 polymerase were seeded onto 6-well plates at 3 × 10^5^ cell per well in duplicate. The next day, cells were transfected with 2 μg of pNDV-HXP-S, 1 μg of pTM1-NP, 0.5 μg of pTM1-P, 0.5 μg of pTM1-L and 1 μg of pCI-T7opt and were re-suspended in 250 μl of a modified Eagle’s Minimum Essential Medium (Opti-MEM; Gibco). The plasmid cocktail was then gently mixed with 15 μL of TransIT LT1 transfection reagent (Mirus, GA, USA). The growth media was replaced with Opti-MEM during transfection. To increase rescue efficiency, BSRT7-DF-1 co-culture was established the next day as described previously [24]. Specifically, transfected BSRT7 and DF-1 cells were washed with warm PBS and trypsinized. Trypsinized cells were neutralized with excessive amount of growth media. BSRT7 cells were mixed with DF-1 cells (~1: 2.5) in a 10-cm dish. The co-culture was incubated at 37°C overnight. The next day, the media was removed, and cells were gently washed with warm PBS, Opti-MEM supplemented with 1% P/S and 0.1 μg/ml of (tosyl phenylalanyl chloromethyl ketone) TPCK-treated trypsin was added. The co-cultures were incubated for 2 or 3 days before inoculation into 8- or 9-day-old specific pathogen free (SPF) embryonated chicken eggs (Charles River Laboratories, CT, USA). To inoculate eggs, cells and supernatants were harvested and homogenized by several syringe strokes. One or two hundred microliters of the mixture were injected into each egg. Eggs were incubated at 37 °C for 3-5 days and cooled at 4°C overnight. Allantoic fluids (AF) were harvested from cooled eggs and the rescue of the viruses was determined by hemagglutination (HA) assays. RNA of the rescued virus was extracted, and RT-PCR was performed to amplify the cDNA segments of the viral genome. The cDNA segments were then sequenced by Sanger sequencing (Psomagen, MA, USA). The genetic stability of the recombinant viruses was evaluated across multiples passages in 10 days old-SPF embryonated chicken eggs.

### Virus titration by EID_50_ assays

Fifty percent of (embryonated) egg infectious dose (EID_50_) assay was performed in 9 to 11-day old chicken embryonated eggs. Virus in allantoic fluid was 10-fold serially diluted in PBS, resulting in 10^-5^ to 10^-10^ dilutions of the virus. One hundred microliters of each dilution were injected into each egg for a total of 5-10 egg per dilution. The eggs were incubated at 37 °C for 3 days and then cooled at 4°C overnight, allantoic fluids were collected and analyzed by HA assay. The EID_50_ titer of the NDV, determined by the number of HA-positive and HA-negative eggs in each dilution, was calculated using the Reed and Muench method.

### Preparation of inactivated concentrated virus

The viruses in the allantoic fluid were first inactivated using 0.05% beta-propiolactone (BPL) as described previously [23]. To concentrate the viruses, allantoic fluids were clarified by centrifugation at 4,000 rpm at 4°C for 30 min using a Sorvall Legend RT Plus Refrigerated Benchtop Centrifuge (Thermo Fisher Scientific). Clarified allantoic fluids were laid on top of a 20% sucrose cushion in PBS (Gibco). Ultracentrifugation in a Beckman L7-65 ultracentrifuge at 25,000 rpm for 2 hours at 4°C using a Beckman SW28 rotor (Beckman Coulter, CA, USA) was performed to pellet the viruses through the sucrose cushion while soluble egg proteins were removed. The virus pellets were re-suspended in PBS (pH 7.4). The total protein content was determined using the bicinchoninic acid (BCA) assay (Thermo Fisher Scientific).

### SDS-PAGE and Western Blot

The concentrated NDV-HXP-S or WT NDV was mixed with Novex™ Tris-Glycine SDS Sample Buffer (2X) (Thermo Fisher Scientific), NuPAGE™ Sample Reducing Agent (10 x) (Thermo Fisher Scientific) and PBS at appropriate amounts to reach a total protein content. The mixture was heated at 90 °C for 5 min. The samples were mixed by pipetting and loaded to a 4-20% 10-well Mini-PROTEAN TGXTM precast gel. Ten microliters of the Novex™ Sharp Pre-stained Protein standard (Thermo Fisher Scientific) was used as the ladder. The electrophoresis was run in Tris/Glycine SDS/Buffer (Bio-Rad).

For comassie blue staining, the gel was washed with distilled water at room temperature several times until the dye front in the gel was no longer visible. The gel was stained with 20 mL of SimplyBlue™ SafeStain (Thermo Fisher Scientific) for a minimum of 1 h to overnight. The SimplyBlue™ SafeStain was decanted and the gel was washed with distilled water several times until the background was clear. Gels were imaged using the Bio-Rad Universal Hood IiIMolecular imager (Bio-Rad) and processed by Image Lab Software (Bio-Rad).

For Western Blot, proteins were transferred onto polyvinylidene difluoride (PVDF) membrane (GE Healthcare, IL, USA). The membrane was blocked with 5% dry milk in PBS containing 0.1% v/v Tween 20 (PBST) for 1h at RT. The membrane was washed with PBST on a shaker 3 times (10 min at RT each time) and incubated with primary antibodies diluted in PBST containing 1% bovine serum albumin (BSA) overnight at 4°C. To detect the spike protein of SARS-CoV-2, a mouse monoclonal antibody 2B3E5 recognizing the S1 kindly provided by Dr. Thomas Moran at ISMMS was used. The HN protein was detected by a mouse monoclonal antibody 8H2 (MCA2822, Bio-Rad). The membranes were then washed with PBST on a shaker 3 times (10 min at RT each time) and incubated with sheep anti-mouse IgG linked with horseradish peroxidase (HRP) diluted (1:2,000) in PBST containing 5% dry milk for 1h at RT. The secondary antibody was discarded and the membranes were washed with PBST on a shaker 3 times (10 min at RT each time). Pierce™ ECL Western Blotting Substrate (Thermo Fisher Scientific) was added to the membrane, the blots were imaged using the Bio-Rad Universal Hood II Molecular imager (Bio-Rad) and processed by Image Lab Software (Bio-Rad).

### Animal experiments

All the animal experiments were performed in accordance with protocols approved by the Icahn School of Medicine at Mount Sinai (ISMMS) Institutional Animal Care and Use Committee (IACUC). All experiments with live SARS-CoV-2 were performed in the Centers for Disease Control and Prevention (CDC)/US Department of Agriculture (USDA)-approved biosafety level 3 (BSL-3) biocontainment facility of the Global Health and Emerging Pathogens Institute at the Icahn School of Medicine at Mount Sinai, in accordance with institutional biosafety requirements.

### Mouse immunization and challenge studies

Female BALB/c mice were used in all studies. Intramuscular vaccination using 1 μg of total protein of inactivated NDV-HXP-S vaccine or negative control WT NDV was prepared in 100 μl total volume. Two immunizations were performed for all the mice with a 21-day interval. For SARS-CoV-2 infection, mice were intranasally infected with 2.5 × 10^8^ plaque forming units (PFU) of Ad5-hACE2 5 days prior to being challenged with 1 × 10^5^ PFU USA-WA1/2020 strain (Wuhan-like), 3.4 × 10^4^ PFU of the hCoV-19/USA/MD-HP01542/2021 JHU strain (Beta, kindly provided by Dr. Andrew Pekosz from Johns Hopkins Bloomberg School of Public Health), 6.3 × 10^4^ PFU of the hCoV-19/Japan/TY7-503/2021 strain (Gamma), 1.6 × 10^5^ PFU of the hCoV-19/USA/NYMSHSPSP-PV29995/20212021 strain (Delta, obtained from Dr. Viviana Simon (Mount Sinai Pathogen Surveillance program) and 5.0 × 10^3^ PFU of the hCoV-19/USA/WI-UW-4340/2021strain2021strain (Mu, kindly provided by Dr. Michael S. Diamond from Washington University Medical School in St. Louis). Viral titers in the lung homogenates of mice 2 days post-infection were used as the readout for protection. Briefly, the lung lobes were harvested from a subset of animals per group and homogenized in 1 mL of sterile PBS. Viral titers in the lung homogenates were measured by plaque assay on Vero-E6 or Vero-TMPRSS2 cells. Blood was collected by submandibular vein bleeding. Sera were isolated by low-speed centrifugation and stored at −80°C before use.

### Recombinant proteins

Recombinant WA1, Beta, Alpha and Omicron spike proteins and recombinant WA1, Beta, Alpha, Gamma, Delta and Omicron RBDs were generated and expressed in Expi293F cells (Life Technologies, Thermo Fisher Scientific) as previously described [25, 26]. Proteins were then purified after transient transfections with each respective plasmid. Briefly, the mammalian-cell codon-optimized nucleotide sequence of a soluble spike protein (amino acids 1-1,213) lacking the polybasic cleavage site, carrying two stabilizing mutations (K986P and V987P), a signal peptide, and at the C-terminus a thrombin cleavage site, a T4 fold-on trimerization domain, and a hexahistidine tag was cloned into the mammalian expression vector pCAGGS. https://www.beiresources.org/).Protein was purified using gravity flow purification with Ni-nitrilotriacetic acid (NTA) agarose (Qiagen, Germany) and concentrated and buffer exchanged in Amicon centrifugal units (EMD Millipore, MA, USA). The purified recombinant proteins were analyzed via reducing sodium dodecyl sulfate-polyacrylamide gel electrophoresis (SDS-PAGE). The desired protein folding was confirmed through ELISAs using the Receptor Binding Domain (RBD)-specific monoclonal antibody CR3022 [27]. Recombinant Gamma (10795-CV-100) and Delta (10878-CV-100) spike proteins were purchased from R&D Systems (R&D Systems, Bio-Techne, MN, USA).

### Enzyme-linked immunosorbent assay (ELISA)

Spike-specific IgG in mice sera vaccinated with NDV-HXP-S was measured by ELISA as described previously [23, 28]. Proteins were coated onto Immulon® 4 HBX 96-well microtiter plates (Thermo Fisher Scientific) at 2 μg/mL in 1x coating buffer (SeraCare Life Sciences Inc., MA, USA) at 50 μL/well overnight at 4°C. All plates were washed 3 times with 225 μL PBS containing 0.1% (vol/vol) Tween-20 (PBST) and 220 μL blocking solution (3% goat serum, 0.5% non-fat dried milk powder, 96.5% PBST) was added to each well and incubated for 1 hour at RT. Individual serum samples or pooled sera were serially diluted 3-fold in blocking solution followed by a 2-hour incubation at RT at a starting dilution of 1:30. ELISA plates were afterwards washed 3 times with PBST and 50 μL of anti-mouse IgG-horseradish peroxidase (HRP) conjugated antibody (Cytiva, GE Healthcare) was added at a dilution of 1:3,000 in blocking solution. After 1 hour, plates were washed 3 times with PBST and developed using SigmaFast OPD (Sigma-Aldrich, MI, USA) for 10 minutes. Reactions were stopped by adding 50 μL 3M hydrochloric acid and absorbance at 492 nm was determined on a Synergy 4 plate reader (BioTek, Agilent Technologies inc., CA, USA) or similar. For each ELISA plate, the blank average absorbance plus 3 standard deviations was used as a cutoff to determine endpoint titers and the area under the curve (AUC) using GraphPad Prism.

### Microneutralization assays using the authentic SARS-CoV-2 viruses

Microneutralization assays using the authentic SARS-CoV-2 viruses were performed as described previously in Vero-TMPRSS2 [5]. All procedures were performed in a BSL-3 facility at the Icahn School of Medicine at Mount Sinai following standard safety guidelines. Vero-TMPRSS2 cells were seeded in 96-well high binding cell culture plates (Costar, Corning) at a density of 20,000 cells/well in complete Dulbecco’s modified Eagle medium (cDMEM) one day prior to the infection. Heat inactivated serum samples (56°C for 1 hour) were serially diluted (3-fold) in minimum essential media (MEM; Gibco) supplemented with 2 mM L-glutamine (Gibco), 0.1% sodium bicarbonate (w/v, HyClone), 10 mM HEPES (Gibco), 100 U/ml penicillin, 100 μg/ml streptomycin (P/S; Gibco) and 0.2% BSA (MP Biomedicals, CA, USA) starting at 1:10. Remdesivir (Medkoo Bioscience inc., NC, USA) was included to monitor assay variation. Serially diluted sera were incubated with 10,000 TCID50 per mL of Wuhan-like WT USA-WA1/2020 SARS-CoV-2, PV29995/2021 (B.1617.2, Delta), MSHSPSP-PV27007/2021 (B.1.351, Beta) and PV44488/2021 (B.1.1.529, Omicron) for one hour at RT, followed by the transfer of 120 μl of the virus-sera mix to Vero-TMPRSS2 plates. Infection proceeded for one hour at 37°C and inoculum was removed. 100 μl/well of the corresponding antibody dilutions plus 100 μl/well of infection media supplemented with 2% FBS (Gibco) were added to the cells. Plates were incubated for 48h at 37°C followed by fixation overnight at 4°C in 200 μl/well of a 10% formaldehyde solution. For staining of the nucleoprotein, formaldehyde solution was removed, and cells were washed with PBS (pH 7.4) (Gibco) and permeabilized by adding 150 μl/well of PBS, 0.1% Triton X-100 (Fisher Bioreagents, MA, USA) for 15 min at RT. Permeabilization solution was removed, plates were washed with 200 μl/well of PBS (Gibco) twice and blocked with PBS, 3% BSA for 1 hour at RT. During this time the primary antibody was biotinylated according to manufacturer protocol (Thermo Scientific EZ-Link NHS-PEG4-Biotin). Blocking solution was removed and 100 μl/well of biotinylated mAb 1C7C7, a mouse anti-SARS nucleoprotein monoclonal antibody generated at the Center for Therapeutic Antibody Development at The Icahn School of Medicine at Mount Sinai ISMMS (Millipore Sigma) at a concentration of 1μg/ml in PBS, 1% BSA was added for 1 hour at RT. Cells were washed with 200 μl/well of PBS twice and 100 μl/well of HRP-conjugated streptavidin (Thermo Fisher Scientific) diluted in PBS, 1% BSA were added at a 1:2,000 dilution for 1 hour at RT. Cells were washed twice with PBS, and 100 μl/well of Sigmafast OPD (Sigma-Aldrich) were added for 10 min at RT, followed by addition of 50 μl/well of a 3 M HCl solution (Thermo Fisher Scientific). Optical density (OD) was measured (490 nm) using a microplate reader (Synergy H1; Biotek). Analysis was performed using GraphPad Prism 7 software. After subtraction of background and calculation of the percentage of neutralization with respect to the “virus only” control, a nonlinear regression curve fit analysis was performed to calculate the 50% inhibitory dilution (ID50), with top and bottom constraints set to 100% and 0% respectively. All samples were analyzed in a blinded manner.

### SARS-CoV-2 plaque assay

Plaque assays with SARS-Cov-2 viruses were performed in the BSL3 facility. Vero-E6 cells or Vero-TMPRSS2 were seeded onto 12-well plates in growth media at 1:5 and cultured for two days. Tissue homogenates were 10-fold serially diluted in infection medium (DMEM containing 2% FBS, 100 unit/mL of penicillin, 100 μg/mL of streptomycin (P/S; Gibco) and 10 mM HEPES). Two hundred microliters of each dilution were inoculated onto each well starting with a 1:10 dilution of the sample. The plates were incubated at 37°C for 1 h with occasional rocking every 10 min. The inoculum in each well was then removed and 1 mL of agar overlay containing 0.7% of agar in 2 × MEM was placed onto each well. Once the agar was solidified, the plates were incubated at 37 °C with 5% CO_2_. Two days later, the plates were fixed with 5% formaldehyde in PBS overnight before being taken out from BSL3 for subsequent staining under BSL2 conditions. The plaques were immuno-stained with an anti-SARS-CoV-2 NP primary mouse monoclonal antibody 1C7C7 kindly provided by Dr. Thomas Moran at ISMMS. An HRP-conjugated goat anti-mouse secondary antibody was used at 1:2000 and the plaques were visualized using TrueBlue™ Peroxidase Substrate (SeraCare Life Sciences Inc.)

### Phylogenetic tree

The phylogenetic tree was built from 3057 SARS-CoV-2 genomes samples between December 2019 and February 2022. Phylogenetic tree and frequency timelines were obtained from the Nextstrain/ncov Project within the GISAID initiative (www.nextstrain.org/ncov) under CC-BY[29, 30].

### Statistics

A one-way ANOVA with Dunnet multiple comparisons test was used to compare the plaque assay and ELISA biding titers. Statistical analyses were performed using Prism software (GraphPad).

